# The Junction Usage Model (JUM): A method for comprehensive annotation-free analysis of alternative pre-mRNA splicing patterns

**DOI:** 10.1101/116863

**Authors:** Qingqing Wang, Donald C. Rio

## Abstract

Alternative pre-mRNA splicing (AS) greatly diversifies metazoan transcriptomes and proteomes and is crucial for gene regulation. Current computational analysis methods of AS from Illumina RNA-seq data rely on pre-annotated libraries of known spliced transcripts, which hinders AS analysis with poorly annotated genomes and can further mask unknown AS patterns. To address this critical bioinformatics problem, we developed a method called the Junction Usage Model (JUM) that uses a bottom-up approach to identify, analyze and quantitate global AS profiles without any prior transcriptome annotations. JUM accurately reports global AS changes in terms of the five conventional AS patterns and an additional “Composite” category composed of inseparable combinations of conventional patterns. JUM stringently classifies the difficult and disease-relevant pattern of intron retention, reducing the false positive rate of IR detection commonly seen in other annotation-based methods to near negligible rates. When analyzing AS in RNA-samples derived from Drosophila heads, human tumors and human cell lines bearing cancer-associated splicing factor mutations, JUM consistently identified ~ twice the number of novel AS events missed by other methods. Computational simulations showed JUM exhibits a 1.2-4.8 times higher true positive rate at a fixed cut-off of 5% false discovery rate. In summary, JUM provides a new framework and improved method that removes the necessity for transcriptome annotations and enables the detection, analysis and quantification of AS patterns in complex metazoan transcriptomes with superior accuracy.

## INTRODUCTION

Alternative pre-mRNA splicing (AS) is a major gene regulatory mechanism that greatly expands proteomic diversity and serves as a crucial determinant of cell fate and identity. More than 95% of human gene transcripts undergo AS that enables one single gene locus to produce multiple, and usually functionally distinct pre-mRNA and protein isoforms (1, 2). AS is regulated by a large constellation of RNA-binding proteins that interact with cis-acting RNA elements embedded in nuclear pre-mRNA sequences (3, 4). Distinct cellular states or tissue types are associated with different AS profiles that affect almost every aspect of cellular functions, including proliferation, differentiation, apoptosis and migration (1, 5, 6). Furthermore, mutations that result in aberrant AS patterns are a major source for human diseases, such as cancer as well as immune and neurological disorders (7–9). Thus, a thorough and comprehensive evaluation of global AS profiles in different tissues, cells and diseased states will be critical to understand the role of AS in gene regulation and facilitate the development of screening and therapeutic strategies to diagnose, treat and prevent many diseases linked to defects in AS. However, due to the exceptionally diverse and dynamic feature of AS patterns, systematic quantification and analysis of cellular AS profiles among a complex array of tissues or cell types remains a major unsolved challenge in the bioinformatics of gene expression.

Recent technical advances in short-read high-throughput Illumina transcriptome sequencing (RNA-seq) provide powerful tools to investigate AS at the genome-wide scale, but at the same time presents a formidable computational challenge to accurately classify and quantitate global AS changes from raw RNA-seq data. Previously, a number of computational software tools and algorithms have been developed for this purpose (7, 10–22), but most use a top-down approach that relies on pre-annotation of known AS events or an incomplete, pre-annotated transcriptome to draft the general picture of global AS patterns for quantification and analysis. As complete dependency on annotation (10) restricts AS analysis to only previously observed AS events, recent methods generally use two approaches to extend the analysis to unannotated splicing events: 1) supplement the pre-annotated AS event library with novel splice junction-implicated AS events identified from the sample under analysis (15, 18); or 2) provide a *de novo* transcriptome annotation through *ab initio* transcriptome assembly from RNA-seq data using probabilistic models (12, 23–26). For the first approach, the library of pre-annotated AS events is still the primary source for calling AS events and can either mask or misclassify novel AS events in the specific RNA-seq sample. For the second approach, a precise and deterministic *ab initio* assembly of transcriptomes from shotgun RNA sequencing is still a big computational challenge for the field, especially for genes that produce multiple transcripts with complex AS patterns. Thus, the difficulties in transcriptome assembly will directly affect the quality of downstream AS analysis. Considering the caveats described above, there is an urgent need for development of computational tools that can perform accurate, comprehensive, and tissue-specific global AS analysis with a different approach.

Here, we present a new computational method called the Junction Usage Model (JUM) that uses a bottom-up approach to profile, analyze and quantitate tissue-specific global AS patterns without any prior knowledge of the transcriptome. JUM exclusively uses sequence reads spanning splice junctions to faithfully assemble complete AS patterns in the RNA-seq samples based on their unique topological features and to quantify splicing changes. We applied JUM to analyze AS patterns in RNA samples from Drosophila heads, mouse embryonic neurons, human cancer tumor samples, as well as human cell lines bearing cancer-associated splicing factor mutations. We demonstrate that JUM consistently identified numerous novel, previously not observed, true tissue-specific AS events that were missed or misclassified when analyzed using annotation-based methods. Furthermore, computational simulations showed that JUM exhibits superior performance in terms of both specificity and sensitivity compared to several popular annotation-based methods. Thus, JUM provides a new framework and improved analytical approach to study the extraordinarily diverse global cellular AS transcriptome profiles and the dynamic regulation of AS without the necessity of transcriptome annotation. JUM can be readily applied to a wide range of RNA samples from different organisms for accurate and quantitative analysis of differential AS patterns.

## RESULTS

### JUM utilizes sequence reads spanning splice junctions to construct AS structures as the basic quantitation unit for AS analysis

JUM exclusively uses sequence reads that map over splice junctions to detect and quantitate splicing events (Figure 1A), as these reads provide the most direct evidence for the splicing of the corresponding intron and quantitatively reflect the level of splicing. These splice junction reads can be inferred through the mapping of the shot-gun sequencing reads to the genome as reads that cannot be completely mapped to one location in the genome, but instead map as “split” reads. From there, JUM defines the AS structure as the basic quantitation unit for AS analysis. An AS structure is a set of splice junctions that share the same start site or the same ending site, with each splice junction in an AS structure defined as a sub-AS-junction (Figure 1A, Figure 1B). JUM uses AS structures for AS analysis because AS structures are not only the basic graphical nodes that compose the conventionally recognized AS patterns (alternative 5’ splice site – A5SS, alternative 3’ splice site – A3SS, skipped cassette exon – SE, mutually exclusive exons – MXE and intron retention – IR), but also the relative levels of sub-AS-junctions within an AS structure directly reflect the level of alternative splicing, greatly facilitating AS quantification. As a result, an A5SS or A3SS event is composed of one AS structure with two sub-AS-junctions (Figure 1B, Figure 1C); a SE event is composed of two AS structures, each with two sub-AS-junctions (Figure 1D); a MXE event with two mutually exclusive exons is composed of two AS structures, each with two sub-AS-junctions (Figure 1E).

**Figure 1.**
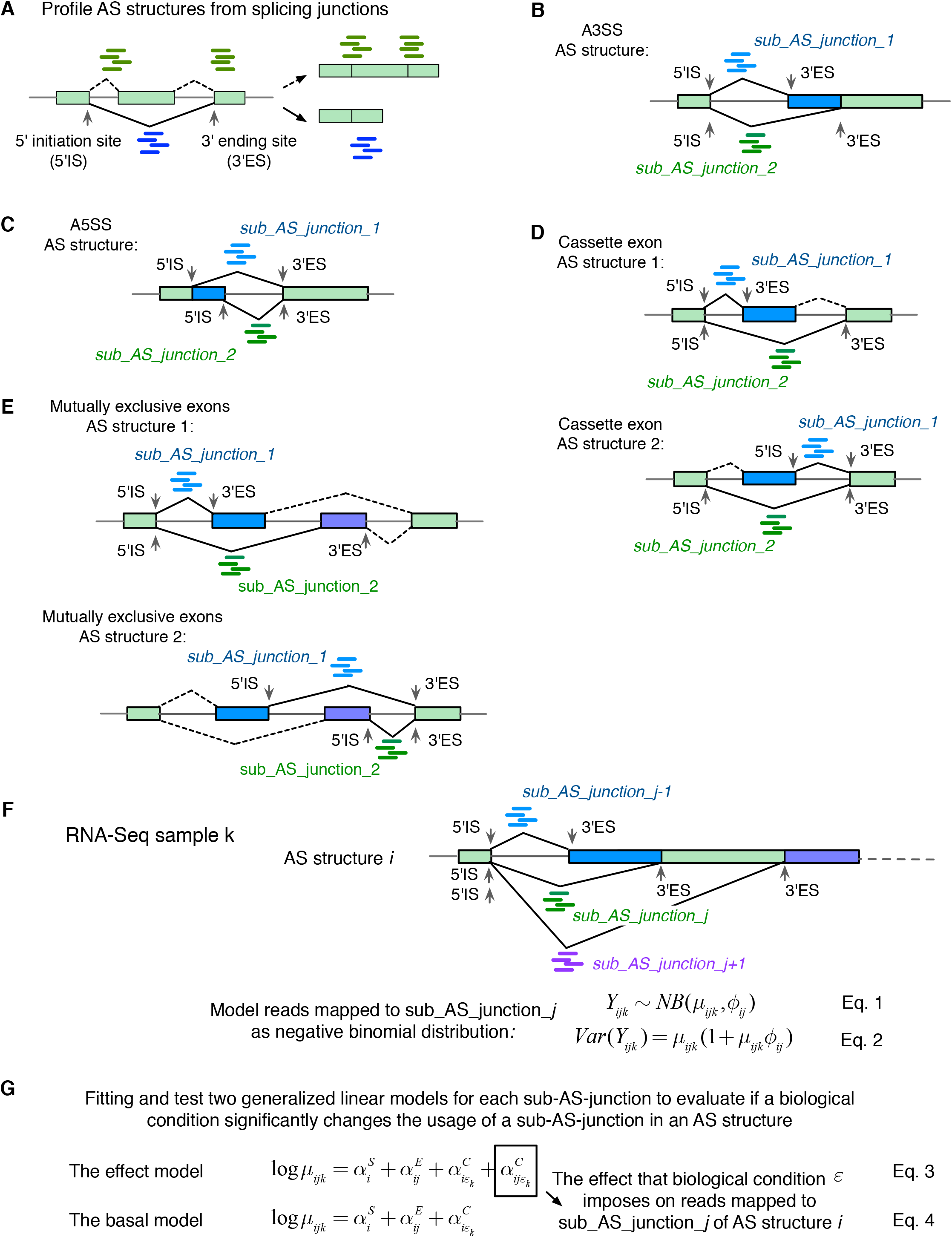
JUM exclusively uses sequence reads mapped to splice junctions and defines AS structures as the basic quantitation unit for differential AS analysis. (A) JUM uses RNA-seq reads mapped to splice junctions for AS quantification. Green rectangles indicate exons and lines introns. Green and blue short lines represent reads that mapped to splice junctions connecting exons, which are the most direct evidence for the existence and quantitative assessment of a given splice junction. JUM defines the start coordinate of a splice junction as the 5’ initiation site (5’IS) and the end coordinate of a splice junction as the 3’ ending site (3’ES). An “AS structure” is defined as a set of junctions that share the same 5’IS or the same 3’ES. Each splice junction in an AS structure is defined as a sub-AS-junction. (B-E) AS structures are the basic element that comprise all conventionally recognized AS patterns. (F) JUM models the sequence reads that map to a sub-AS-junction as negative binomial distribution to quantify the “usage” of each sub-AS-junction in an AS structure under one biological condition. (G) JUM fits two generalized linear models to evaluate the influence of a given biological condition on the usage of a specific sub-AS-junction in an AS structure.

After the profiling of all AS structures, JUM counts sequence reads that are mapped to each sub-AS-junction in every AS structure under a biological condition and defines the read count as the “usage” of a sub-AS-junction relative to other sub-AS-junctions in the same AS structure under that condition. To quantify AS changes, JUM compares the usage of every profiled sub-AS-junction in the AS structure between conditions and profiles for AS structures that contain sub-AS-junctions with differential usage (Figure 1F). To do this, JUM models the total number of reads that map to a sub-AS-junction as negative binomial distribution (Figure 1F, Eq. 1). Negative binomial distributions have been widely applied in high-throughput sequencing data analysis to model read counts, as these models nicely depict the over-dispersion phenomenon observed in next-generation RNA sequencing experiments (11, 27–30). In negative binomial distributions, the variance among biological replicates is dependent on the mean through a parameter that describes dispersion (Figure 1F, Eq. 2). To infer the dispersion parameter, JUM applies a similar empirical Bayesian approach as described (28–31). JUM first estimates a dispersion parameter for each sub-AS-junction with Cox-Reid-adjusted maximum likelihood. JUM then fits a mean-variance function for all sub-AS-junctions from all AS structures on their average normalized count values. Finally, JUM shrinks the dispersion parameter for each individual sub-AS-junction towards the fitted value depending on how close the real dispersion tends to be to the fitted value and the replicate sample size (28–31). To evaluate if a biological condition significantly changes the usage of a sub-AS-junction in the AS structure, JUM adapts a generalized linear model (GLM) approach as described (11, 30, 32), so that two GLM models are fitted and tested for each sub-AS-junction in the AS structure (11) (Figure 1G). The basal model evaluates the effect from the following three elements to the usage of the sub-AS-junction: the basal expression level of the AS structure of the corresponding gene (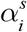, Figure 1G, Eq. 4), the fraction of sequence reads that mapped to each sub-AS-junction from the total number of reads mapped to the AS structure (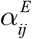, Figure 1G, Eq. 4), as well as the overall change of basal expression of the AS structure upon a biological condition (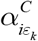, Figure 1G, Eq. 4). On the other hand, the effect model evaluates an additional influence imposed on the usage of a sub-AS-junction by a biological condition (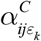, Figure 1G, Eq. 3). The fitting of the effect and basal model are compared and a *χ*^2^ likelihood-ratio test performed (11) so as to test if a biological condition causes significant differential usage of a sub-AS-junction in the AS structure.

### JUM profiles a tissue-specific global AS atlas by faithfully assembling AS structures into conventionally recognized AS patterns without any prior knowledge of the transcriptome annotation

After differential AS analysis using AS structures, JUM assembles profiled AS structures into conventionally recognized categories of AS patterns using graph theory, based on the unique topological feature of each pattern. To do this, JUM first converts each AS pattern into a graph by converting exons into nodes and splice junctions as arcs that connect exon nodes. JUM then defines a frequency parameter *S_I_* for each sub-AS-junction as the number of AS structures that share the specific sub-AS-junction. Because of the definition of AS structures, it can be proven that a given sub-AS-junction can only be included in up to two AS structures (i.e. *S_I_* can only be 1 or 2). For the A5SS or A3SS patterns, the representative graphs are asymmetric and are composed of one AS structure with *S_I_* value equal to 1 for all sub-AS-junctions (Figure 2A, Figure 2B). For the SE pattern, the representative graphs are symmetric, composed of two AS structures, each containing two sub-AS-junctions with *S_I_* values equal to 1 and 2, respectively (Figure 2C). For SE, JUM utilizes extra quarantine steps here, including tiled sequence reads that support the coverage over the entire cassette exon region to avoid false positive calls. For MXE with *n* mutually exclusive exons, the representative graph is composed of one pair of AS structures that each has *n* sub-AS-junction with *S_I_* values all equal to 1 (Figure 2D). For the MXE pattern, JUM utilizes extra quality control steps, including that coordinates of MXE exons meet the condition 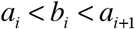, where ^*i* = 1, …*n*^ (Figure 2D) and tiled sequence reads that support coverage over the entire regions of all mutually exclusive exons. Based on the unique topological features of each AS pattern described above, JUM searches for sets of AS structures that match the composition of each AS pattern and bundle them together as one AS event under the corresponding AS pattern category.

**Figure 2.**
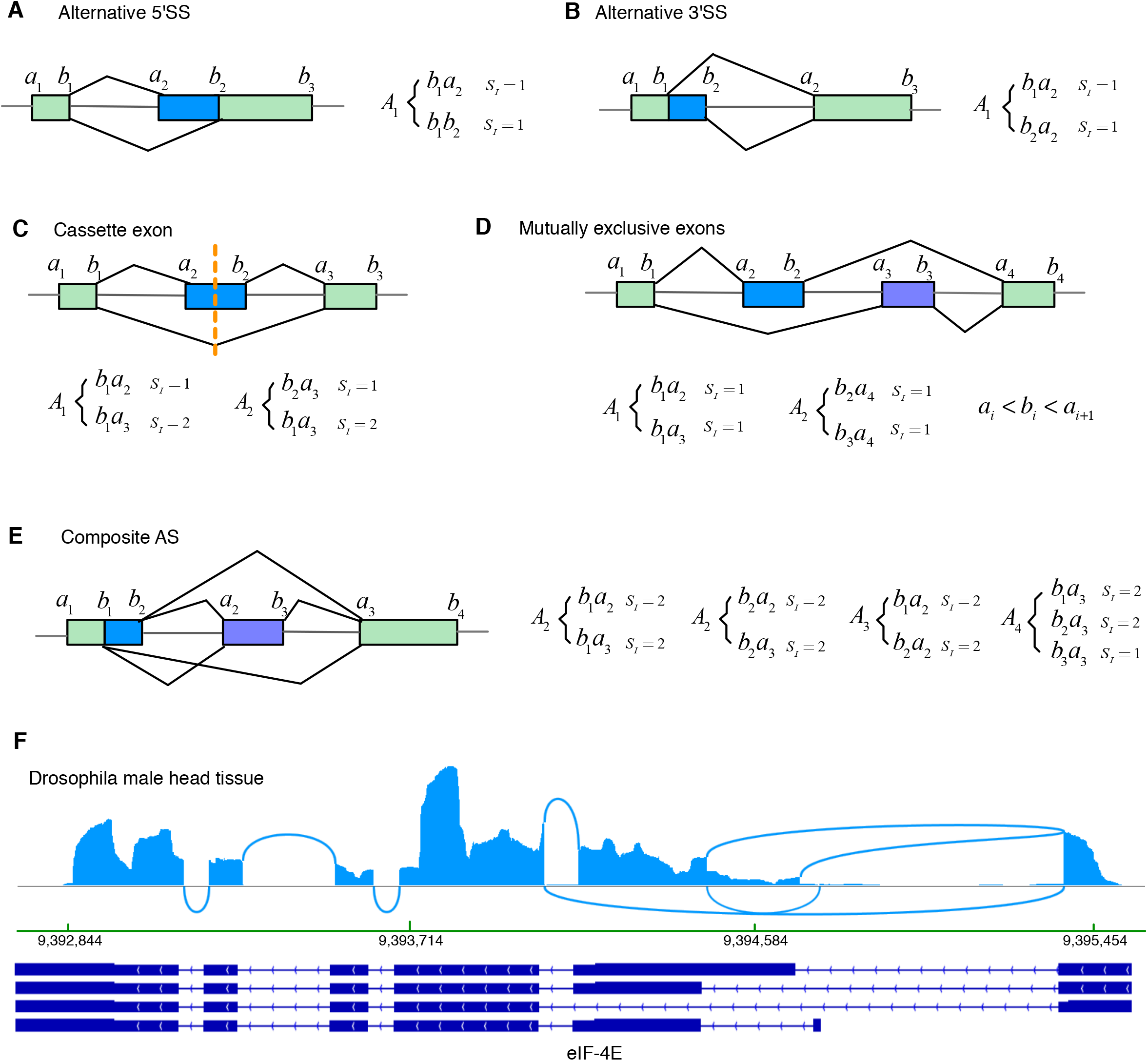
JUM profiles the AS atlas specific to the sample by assembling AS structures into conventionally recognized categories of AS patterns based on the unique topological features of each AS pattern type. A-D) The topological features of AS patterns: A5SS (A), A3SS (B), SE (D), and MXE (D) represented by the splicing graphs that is composed of unique set of AS structures and the values of frequency parameter *S_I_* of each sub-AS-junction in the AS structures. (E) JUM defines an additional, previously unclassified AS pattern category—the “Composite AS” which is a complex combination of several conventionally recognized AS patterns. An example for such a “composite” AS pattern is shown in (F) for the eIF-4E gene transcripts found in Drosophila male head tissue RNA-seq samples (46). Arcs represent splice junctions that connect different exons.

Additionally and uniquely, JUM also recognizes and defines an additional AS pattern called “Composite”, which describes an AS event that is an inseparable combination of several conventionally recognized AS patterns (Figure 2E, Figure 2F). Such composite AS patterns are found extensively in Drosophila, rodent and human tissues and cell line RNA samples (Figure 2F), further illustrating the complexity and diversity of AS in complex tissues. For example, JUM identifies an AS event in the transcripts from the eIF-4E gene in Drosophila male heads that is an intertwined combination of SE, A5SS and A3SS and cannot be de-convoluted into any of the three individual patterns separately (Figure 2F). Although such complex AS events have been reported before (33), they are mostly divided into the conventionally recognized AS patterns by the currently available AS analysis tools. This approach overlooks the fact that such composite AS events are usually an intertwined combination of the basic AS patterns and a forced separation of these events into the basic patterns can lead to inaccurate quantification of the AS changes. JUM thus classifies these complex AS events in their own category. For Composite AS events in JUM, the representative graph includes an interconnected combination of AS structures corresponding to each AS pattern that composes the composite AS pattern. For example, a composite AS pattern event that is a combination of an A5SS and SE is composed of four AS structures, three of which have sub-AS-junctions with *S_I_* values all equal to 2 and one has one sub-AS-junction with an *S_I_* value=1 and the rest with *S_I_* value=2 (Figure 2E).

For each AS event in the corresponding AS pattern category, JUM not only reports the statistical test score for differential splicing, but also a “delta Percent-Spliced-in” (delta_PSI or ΔΨ) measure that reflects the level of splicing changes between conditions. For SE or IR patterns in which each AS event only includes two alternatively spliced isoforms, ΔΨ is calculated as the change in the percentage of the exon-inclusion isoform or the intron-inclusion isoform between conditions, respectively. For A5SS, A3SS, MXE and Composite events however, since each AS event can include multiple AS isoforms, ΔΨ is calculated instead as the change in the “usage” (or, percentage) of each sub-AS-junction in the AS structures that corresponds to each AS isoform.

### JUM applies stringent criteria for the analysis of intron retention (IR) events

Intron retention (IR) has been a relatively under-studied category compared to other AS patterns, but nevertheless a key AS pattern. IR events have been reported to be frequently found in mammals and have been shown to play crucial roles for the normal functioning of the organism and in disease in eukaryotes (34, 35). For example, tissue-specific IR of the Drosophila P element transposase pre-mRNA underlies the restriction of transposon activity to germ line tissues (36, 37). Recently, a bioinformatics study reported that widespread retained introns were associated with various cancer types compared to matched normal tissues (38). Increased IR has also been shown to be associated with the pluripotent state of embryonic stem cells (39). However, intron retention is an intricate AS pattern that can be easily misclassified. The most common approaches to quantify retained intron-containing isoforms in currently available AS analysis tools are either to use the sum of sequence reads mapped to the upstream exon-intron boundary and the downstream intron-exon boundary, or to use any reads mapped to the intronic region or just the center of the intron (Figure 3A). A major caveat of these approaches is that other AS patterns can be mistaken as IR, especially when alternative SE or MXE exons reside within an intron (Figure 3B, Figure 3C, Figure 3D) or an A5SS/A3SS event resides at the edge of the intron (Figure 3D). In such scenarios, sequence reads from the intronic region can in fact come from the SE or MXE exons and reads mapped to exon-intron or intron-exon boundaries can come from A5SS or A3SS events, but with the conventionally available methods these reads can be mistakenly interpreted as support for intron-retained isoforms.

**Figure 3.**
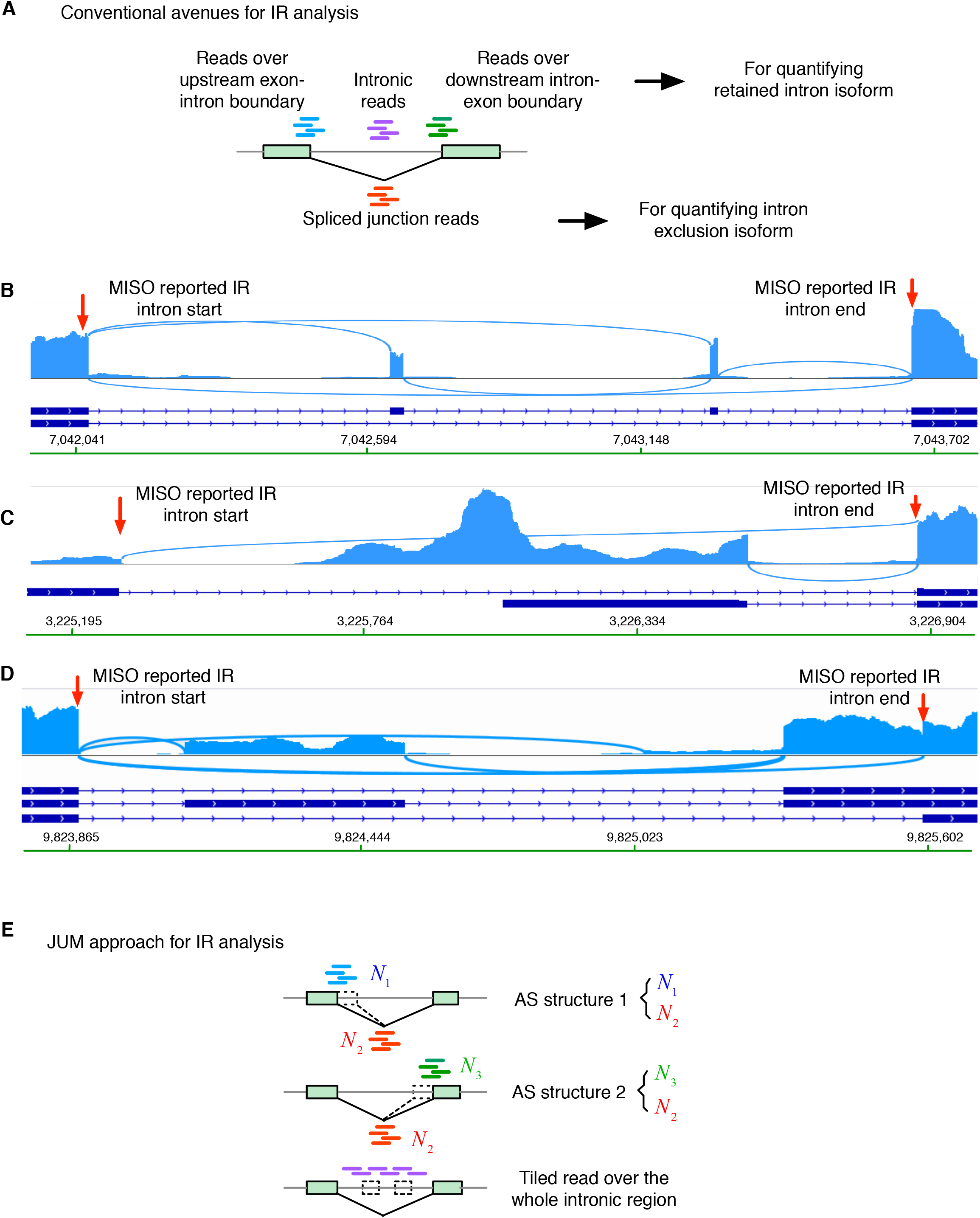
JUM applies stringent criteria for detection, quantitation and analysis of IR events. (A) The conventional avenues for IR analysis in the current available AS analysis tools. RNA-seq reads spanning intron-exon or exon-intron boundaries are represented by short green or blue lines, respectively. Short purple lines represent sequence reads mapped to intronic regions. Short red lines represent sequence reads mapped to the splice junction for the corresponding intron. (B-D) The commonly used strategies in other AS analysis software can mis-classify other AS patterns as IR. Three MISO-reported (46) significantly changed IR events were shown that actually MXE (B), alternative promoter event (C), SE mixed with A3SS (D) from Drosophila male head tissue in a comparison of control wildtype fly strain and a transgenic fly strain that expresses the truncated PSI protein. The start and end points of the retained intron events reported by MISO are denoted by red arrows. (E) The approach that JUM uses to analyze IR. Short blue and green lines represent reads mapped to the exon-intron or intron-exon boundaries, respectively. Short red lines represent sequence reads mapped to the splice junctions. Two AS structures are constructed to analyze the level of retained intron isoform versus spliced intron isoform. Short purple reads represent sequence reads mapped to the intronic regions and are required to be approximately uniformly distributed across the entire intronic region of the retained intron.

To avoid false-positive calls of IR as described above, JUM applies a stringent three-criteria strategy to profile and analyze IR patterns (Figure 3E). First, JUM profiles for splice junctions that do not overlap with any other splice junctions from the RNA-seq data. Second, for each of the resulting splice junctions and the corresponding intron, JUM counts the number of sequence reads mapped to the upstream exon-intron boundary (*N*_1_), reads spanning across the splice junction (*N*_2_), and reads mapped to the downstream intron-exon boundary (*N_3_*), respectively (Figure 3E). JUM then defines two AS structures for each candidate intron: *N* versus *N*_2_, and *N*_3_ versus *N*_2_. Both AS structures must be differentially “used” with the same trend (*N*_1_ and *N*_3_ both significantly more used than *N*_2_ or both significantly less used than *N*_2_) in order for the candidate to be classified as a potential IR event. These two criteria are set to avoid A5SS or A3SS events to be mistaken as IR (Figure 3D). Finally, JUM requires evidence from mapped sequence reads that are approximately evenly distributed across every base of the intron, in order to confirm a real IR event (Figure 3E). This criterion aims at preventing SE or MXE events to be misclassified as IR, as reads from SE or MXE exons residing in the intron will present a much higher, “spikey” distribution pattern compared to other regions of the intron.

### JUM demonstrates superior performance in both specificity and sensitivity compared to other methods in computationally simulated RNA-seq experiments

In order to fully assess the performance of JUM in differential global AS analysis, we performed computational simulations of RNA-seq datasets with varying degrees of alternative splicing in a pre-fixed set of genes (Table S1) and used the simulated datasets to test the ability of JUM to profile true differentially spliced AS events. We compared JUM to five other commonly used annotation-based tools – MISO (10), Whippet (22), Cufflinks (40), MAJIQ (18), and rMATS (15) (Table S2). The five tools were chosen to represent the three most commonly used strategies in annotation-based AS analysis methods: MISO and Whippet depend on either a developer-provided (MISO) or a user-provided (Whippet) pre-annotation of AS events and cannot detect novel AS events outside of the provided database (10, 22); Cufflinks first performs the challenging *de novo* transcriptome assembly from shot-gun sequencing and then quantifies AS changes based on the annotation of the assembled transcriptome (40); MAJIQ and rMATS both use a pre-annotation of the transcriptome to guide the AS analysis, but add in novel splicing junctions detected in the specific RNA-seq sample (15, 18). The latter three methods can extend analysis to novel splicing events in the sample. It should be noted that there are other software tools that have been developed in the completely annotation-dependent category. Here, we chose MISO and Whippet as representatives, as MISO is among the most cited tool in this category and Whippet is among the most recently developed tools in the category (10, 22).

The test datasets are simulated based on real experimental RNA-seq data in Drosophila Schneider-2 cell lines comparing AS changes brought by the RNA interference (RNAi) knockdown of a splicing factor called PSI (41) and a non-targeting, control RNAi knockdown. The simulation follows a method that has been previously described (42). Specifically, a randomly chosen 2,000 genes from expressed genes in the Drosophila Schneider-2 cell line are set to be alternatively spliced, serving as the “true” AS genes, with AS patterns covering all patterns. Triplicates of ~80 million total 100bp RNA-seq reads were simulated for both the PSI knockdown and control knockdown samples. Three independent simulations were performed, with 20%, 40% and 60 % differential splicing changes in the true AS genes between the control and knockdown conditions, respectively. The performance of each AS analysis software under comparison was evaluated in terms of the Receiver Operating characteristic (ROC) curves and the Area Under the Curve (AUC) metric. The ROC curve depicts the true-positive rate (sensitivity) against the false-positive rates (1-specificity) for each threshold setting to call an AS event. AUC is a numerical metric that determines how well a method can distinguish between the true AS events and non-AS events. AUC scores range between 0.5 and 1, and a higher AUC score indicates a method with better discrimination between AS and non-AS events.

Importantly, JUM received the best AUC score in all three simulated RNA-seq experiments among the six methods tested, indicating its superior performance in both sensitivity and specificity (Figure 4A, Figure 4B, Figure 4C). The AUC scores for JUM in all three simulations ranged from 0.92 to 0.95, indicating that JUM is a superior method for accurate differential AS analysis by AUC standard (AUC value between 0.9 and 1). rMATS and MAJIQ show comparable performance, with AUC ranging between 0.83-0.88; Cufflinks performs worse than both rMATS and MAJIQ, with AUC ranging between 0.62-0.66, but still better than the completely annotation-dependent methods Whippet and MISO, which have AUC values of 0.55–0. 64 and 0.52-0.55, respectively (Figure 4A, Figure 4B, Figure 4C). The poor performance of MISO is to some extent understandable, because the true AS genes in the test set used here are randomly chosen, which is vastly different from the developer-provided MISO annotation. The results from these simulations further demonstrate the importance for an AS analysis method to account for sample-specific, novel AS patterns rather than using a pre-annotated library of AS events in order to achieve better performance in AS analysis.

**Figure 4.**
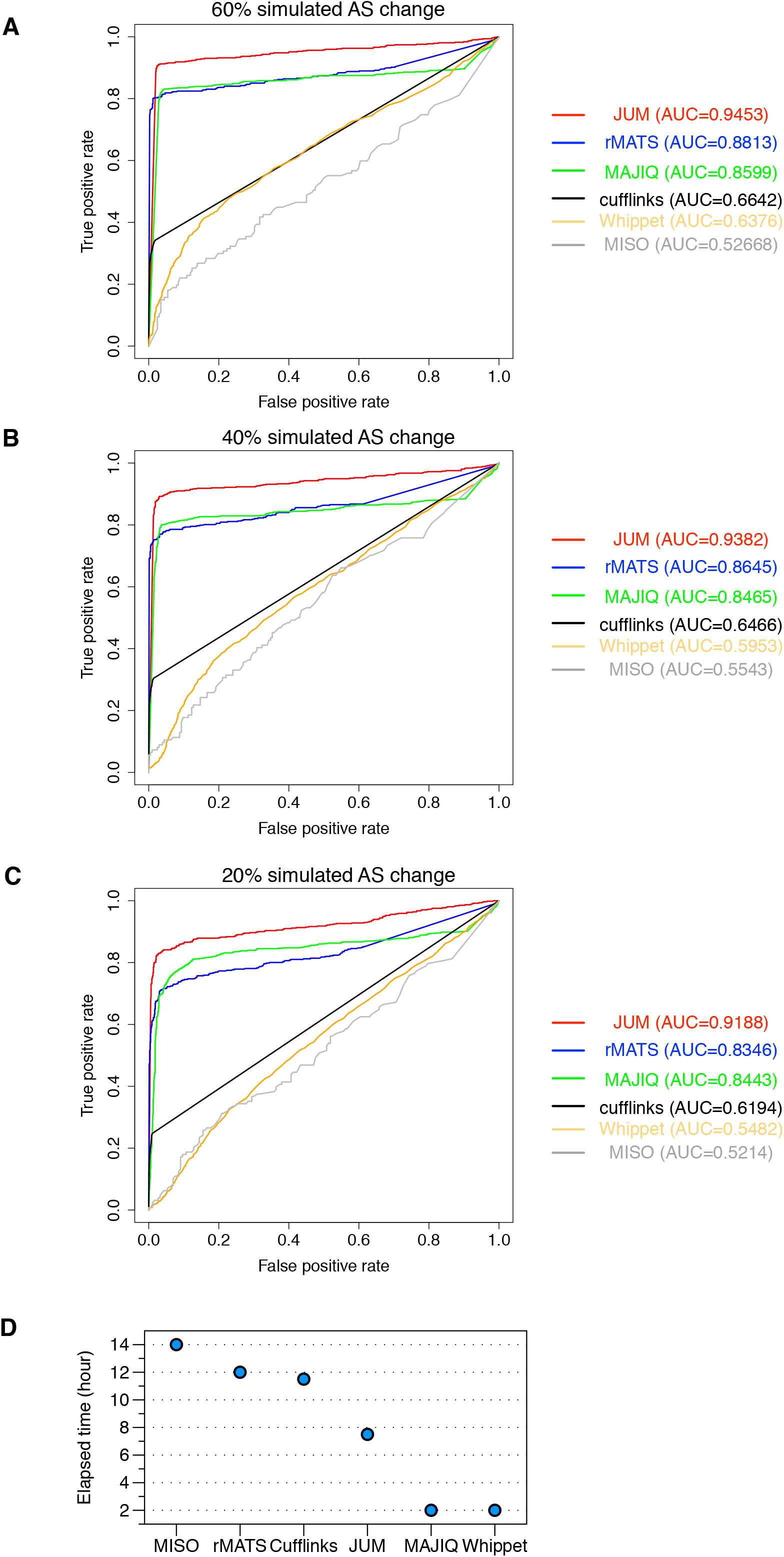
Comparison of JUM to five other most widely used computational methods (rMATS, MAJIQ, cufflinks, MISO and Whippet) for AS analysis using computationally simulated RNA-seq experiments. (A,B,C) ROC (Receiver Operating Characteristic) curves are shown for each method to illustrate their sensitivity and specificity in identifying true differentially spliced AS events. Y axis of the ROC figure shows true positive rate and X axis shows false positive rate. The metric AUC (Area Under an ROC Curve) is listed on the right. The higher the AUC value, the more sensitive and accurate a method is in the performance of differential AS analysis. Among the five methods, MISO and Whippet completely depend on pre-annotated libraries of AS events and cannot detect novel AS events. rMATS, MAJIQ and Cufflinks aid the AS analysis with different workarounds to include novel splicing detection besides the pre-annotated libraries of AS events. JUM performs AS analysis without any prior knowledge of AS. Three independent simulations are done by varying the alternative splicing changes at levels of 20%, 40% and 60%, respectively. (D) The computation time that each software takes to analyze the described simulated data – two sets of triplicates of 80 million 100bp reads are listed.

We also compared the computation time of the six software tools in analyzing the simulated datasets. For analyzing two sets of triplicates of ~80 million 100bp reads on a standard computing cluster without parallel processing, Whippet and MAJIQ are the fastest, taking about 2 hours, followed by JUM that takes about 7.5 hours, Cufflinks 11 hours, rMATS 12 hours and MISO is the slowest that takes about 14 hours (Figure 4D).

Following the simulation, we applied JUM to multiple experimental RNA-seq datasets from two different organisms in order to evaluate the performance of JUM in real RNA-seq datasets.

### JUM significantly reduced the false positive rate of IR detection in colon cancer patient tumor samples and revealed the heterogeneity of IR splicing in cancer

To assess the performance of JUM in IR analysis, we used JUM to analyze differential splicing of IR between the tumor and the matched normal tissue samples from colon cancer patients in The Cancer Genome Atlas (TCGA) database. A previous study (38) used MISO and the MISO built-in pre-annotation of human AS events to compare the AS profiles between patient tumor and matched normal tissues and reported that extensive intron retention is a prevalent feature and was a highly elevated splicing pattern observed in cancer, with colon cancer among the cancer types where this phenomenon was the most obvious (38). We thus compared the performance of JUM with MISO, as well as rMATS, in analyzing the cancer patient datasets. Moreover, since MISO and rMATS are not particularly well tailored to handle IR, we also compared the performance of JUM with IRFinder, a tool specialized in IR analyses (43).

To avoid technical and sampling bias brought about by factors other than the cancerous state into the analysis, we chose samples from two sets of colon cancer patients in TCGA: 1) five male colon cancer patients that are of similar ages (60-68 years old), same colon tumor type (primary tumor), same vital states (alive) and with matched tumor and normal tissue samples sequenced using the same platform; 2) six female colon cancer patients chosen with similar parameters as described above except with a larger age span from 40-85 years old, as there were limited choices in age for female patient samples in the TCGA database (Table S4). For each patient, we applied MISO, rMATS and JUM to compare the global AS changes between the tumor and matched normal tissues, especially in IR events (Table S5, Table S6). As an example, we summarized the results in IR events for a randomly chosen male patient F46704 in Figure 5A. Here, MISO reported a total of 118 differentially spliced IR events, among which only 14 introns were more retained in tumor samples and 104 introns were more retained in the normal tissue; rMATS identified a total of 33 differentially spliced IR events, among which 7 are more retained in the tumor sample; JUM identified a total of 207 differentially spliced IR events, among which 53 are more retained in the tumor sample; IRFinder identified a total of 224 differentially spliced IR events, among which 46 are more retained in the tumor sample. At least in this particular case of patient F46704, the IR analysis resulting from all three methods contradict what was reported previously (38), with more retained introns in the normal tissue instead. This result indicates individual variations in IR pattern changes among cancer patients.

**Figure 5.**
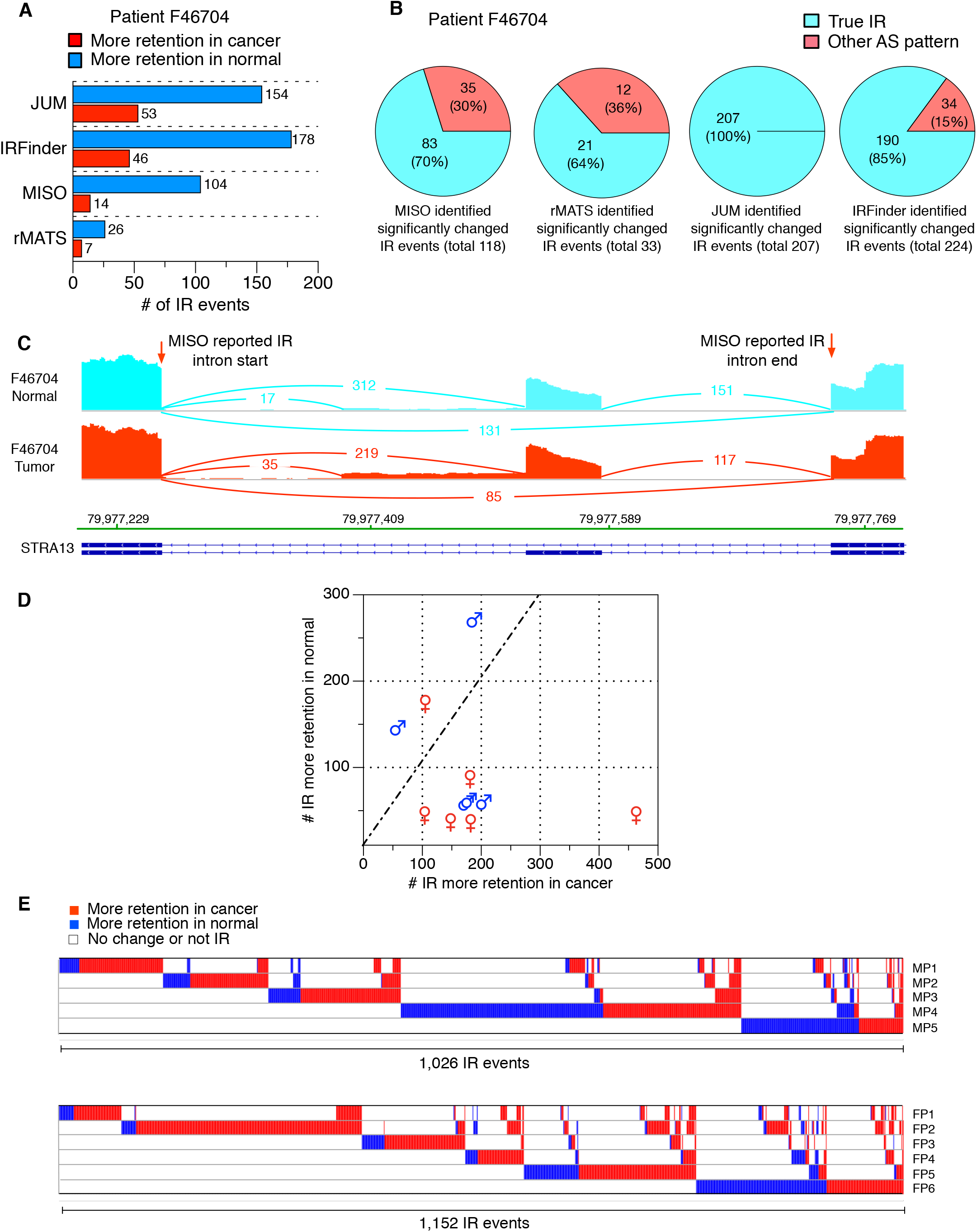
Comparison of JUM, rMATS, MISO and IRFinder in analyzing IR in colon cancer patients tumor versus match normal tissue samples. (A) the number of significantly differentially spliced IR events reported by four methods, both number of IR events that are more retained in cancer and normal tissues are shown for male patient F46704, respectively. (B) Number of false positively reported IR events by four methods for male patient F46704. (C) An example of an incorrectly classified IR event reported by MISO in the gene STRA13 in male patient F46704. The reported intron is specified by red arrows. This MISO-reported “IR” event is in fact a combination of an SE and A5SS event. (D) Distribution of JUM-identified, significantly changed IR events that are more retained in tumor or normal tissues for each male and female patient. (E) Heatmap of all JUM-identified, significantly changed IR events in the set of male patients and female patients, respectively. Every column is an IR event and if the IR event is more retained in the tumor sample, a red grid is shown; if the IR event is more retained in the normal sample, a blue grid is shown. If the IR event did not change between normal or tumor sample or is not an IR event under the patient sample, a white grid is shown.

Considering that IR is an intricate AS pattern that can be easily misclassified, we next examined if the differentially spliced IR events identified by MISO, rMATS, IRFinder and JUM were true IR events. To do this, we chose samples from patient F46704 and visually checked every reported differentially spliced IR event by MISO, rMATS, IRFinder and JUM, respectively, using the genome browser viewer tool igv (44) (Figure 5B). Surprisingly, 30% (35 out of 118) of MISO-reported differentially spliced IR events are not IR events (Figure 5B, Figure 5C). The ratio for rMATS is also high, with 36% (12 out of 33) rMATS-reported IR events not real IR events (Figure 5B, Figure S2A) and IRFinder has much lower false positive rate than MISO and rMATS, with 15% (34 out of 224) of the reported IR events are not real IR events (Figure 5B, Figure S2B). Interestingly, all of the JUM reported IR events are true IR events (Figure 5B). This result indicates that the part of JUM specifically designed for stringent IR analysis significantly reduced the high false positive rate commonly observed in other methods for IR detection.

In order to examine if elevated intron retention is indeed associated with cancer, we further compared the number of more-retained-in-tumor IR events versus more-retained-in-normal-tissue IR events for each of the five male and six female colon cancer patients using JUM (Figure 5D). We found that two out of the five male patients present more retained introns in normal tissues compared to cancer, while the majority (five out of six) of female patients present more retained introns in cancer compared to normal tissues (Figure 5D). This result demonstrates that although elevated intron retention is associated with tumor samples in a subset of colon cancer patients, individual variations among patients are also observed in differential IR splicing events. Importantly, when comparing IR splicing among different patients, we found that these differentially spliced IR events are highly patient-specific, with only two more-retained-in-cancer IR events shared among all five male patients and zero shared among female patients (Figure 5E). The majority of the more-retained-in-cancer IR events identified in one patient either did not exhibit any splicing changes in other patients or were found to be more retained in normal tissues instead, and the same observation is also made for the more-retained-in-normal-tissue IR events. From this analysis, we conclude that alternative splicing of IR events in the colon cancer transcriptomes is highly heterogeneous and specific to individual patients.

### JUM detected significantly more differentially spliced AS events in human cell lines bearing cancer-associated mutation in splicing factor SRSF2 with high accuracy

To further evaluate the sensitivity of JUM in detecting global AS changes in cell samples with complex AS patterns, we used JUM to analyze global AS changes caused by a cancer-associated point mutation (P95H) in the splicing factor SRSF2 in endogenously CRISPR-edited human K562 cell lines (45). Previously, Zhang et al. used rMATS to profile AS changes in the datasets, and reported a total of 548 significantly changed AS events, including 374 SE events, 68 IR events, 15 A3SS events, 25 A5SS events and 66 MXE events (45) (Figure 6A). The number of differentially spliced AS events are distributed with high bias among AS pattern categories, with the majority reported in SE (~68%) and only 5% and 3% in A5SS and A3SS patterns, respectively. By contrast, using JUM with the same statistical cutoff reported in the previous study (45) (adjusted p-value <= 0.1, deltaPSI <= 0.1), we found a total of 1,001 AS events that are differentially spliced in cells carrying the point mutation in SRSF2, almost double the number of events found by rMATS. Among them, JUM found 185 SE events, 135 IR events, 102 A3SS events, 99 A5SS events, 3 MXE events and 477 Composite events, with significantly less bias in the detection of AS events across different AS categories compared to rMATS. Moreover, to test if the distinctively high number of SE events reported by rMATS are real, we visually examined the top 112 most significantly differentially spliced SE events reported by rMATS using igv (44) (Figure 6B). Interestingly, we found about 46% (51 out of 112) of these events are not SE events, but in occur combination with other AS patterns, similar to what JUM classifies as Composite (Figure 6B, Figure 6C). We also examined 112 randomly chosen JUM-reported differentially spliced SE events out of 185 and found 97% of these are indeed true SE events (Figure 6B). In summary, these results demonstrate that JUM can detect significantly more differentially spliced AS event with high accuracy in cell samples with complex AS patterns in comparison to other annotation-based methods like rMATS. A further comparison of the 185 JUM-reported significantly differentially spliced SE events to current human transcriptome annotation revealed that 119 (64%) of these SE were previously known and annotated (Figure 6D) and 66 (36%) correspond to novel cassette exons that are supported by strong evidence from both RNA-seq exon coverage track signals and adjacent splice junctions, through visual examinations of the RNA-seq datasets using igv (44) (Figure 6D, Figure 6B). This observation shows that JUM, although annotation-independent, is capable of accurately profiling AS events that are either previously known or novel to the specific tissue under study.

**Figure 6.**
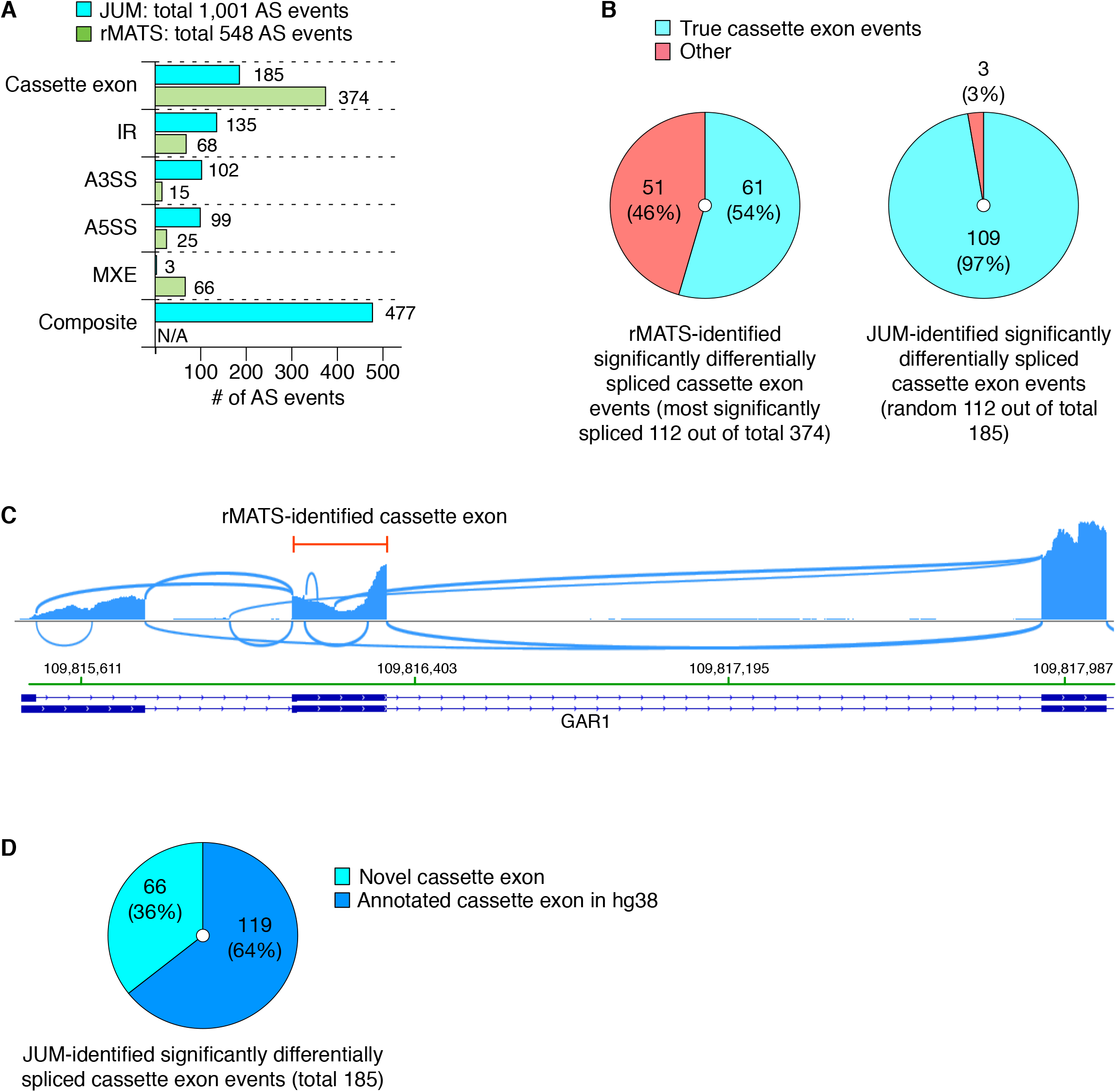
Comparison of JUM and rMATS in analyzing global AS changes brought about by a cancer-associated point mutation in the splicing factor SRSF2 in human K562 cell lines. (A) Number of significantly differentially spliced AS events reported in every AS pattern category by the two methods. (B) Number of cassette exons that are true SE events reported by each method. (C) An example of an incorrectly classified SE event reported by rMATS in the gene GAR1. The reported SE exon is specified by red lines. (D) The distribution of previously annotated and novel SE exons from the 185 JUM-reported significantly changed SE AS events.

### JUM identified significantly more real, novel and functionally important AS events in the head sample of a Drosophila strain carrying a mutation in the splicing factor PSI and is capable of predicting the regulatory function of PSI based on the splicing pattern changes

To assess JUM’s performance in profiling AS changes in tissues with complex AS patterns, we compared the performance of JUM, MISO and rMATS in identifying global AS changes in the male head transcriptome of a Drosophila strain that carries a mutation in the splicing factor PSI leading to the expression of truncated PSI protein (41, 46). The resultant strain exhibits male courtship behavior defects (41). Importantly, the specific mutation in PSI disrupts its interaction with U1 snRNP and thus is expected to affect splicing decisions on a set of target 5’ splice sites (47).

We performed two single-blind, counter tests that compared the performance of JUM to MISO and rMATS. For the first test, we took the set of 21 JUM-identified differentially spliced non-Composite AS events that are functionally linked to the male courtship behavior defects observed in the male Drosophila PSI mutation flies (46), and asked if rMATS and MISO can identify these phenotypically related AS events as well (Figure 7A; Table S8). A visual validation using igv showed that all of these 21 AS events are correctly classified in the corresponding AS pattern category. Among them, we found that the majority of these AS events (12 out of 21, 57%) were identified exclusively by JUM, since neither the annotated AS library for MISO nor the novel splicing-aided mode of rMATS was able to identify these true, novel, and phenotypically-associated AS events in the male PSI mutant fly head samples (Figure 7A, Table S8). Only 1 AS event (5%) was identified by all three methods. Interestingly, this is a SE event which is the most well-annotated AS pattern among all AS pattern categories (Figure 7A, Table S8). 4 AS events are identified by rMATS and JUM but not MISO (19%) and 4 by MISO and JUM but not rMATS (19%) (Figure 7A, Table S8). To confirm that JUM is capable of detecting true AS events that are missed by other software, we performed experimental qRT-PCR validation of the 12 male-courtship-associated AS events that were only identified by JUM but not rMATS or MISO (Figure 7D, Figure S3-S6). Interestingly, all 12 events were validated as true, significantly changed AS events in the PSI mutant male fly head tissue compared to wild type (Figure 7D, Figure S3-S6). These results suggest that JUM is clearly capable of identifying significantly more (in this case two times) functionally-relevant, novel, and tissue-specific AS events that are not recognized by other annotation-based techniques, even when the annotation-based software is aided with a novel splice junction-detection mode.

**Figure 7.**
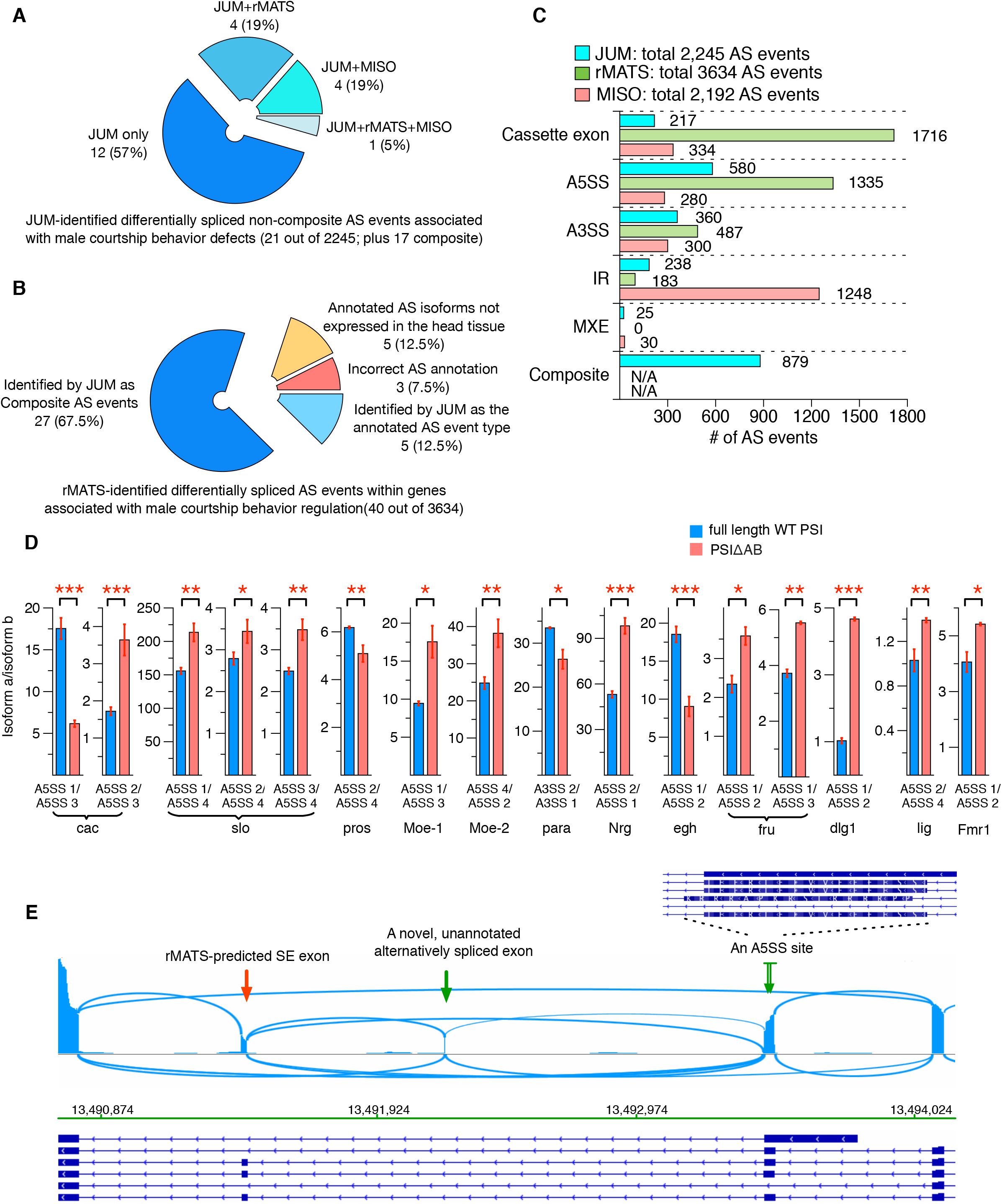
Comparison of JUM, rMATS and MISO in analyzing global AS changes brought about by a truncation mutation in the splicing factor PSI in Drosophila male heads. (A) Test if rMATS and MISO can also identify JUM-predicted, experimentally-validated, and functionally-crucial differentially spliced non-composite AS events that are associated with the male courtship defect phenotype in the male head sample of a PSI mutant Drosophila strain. The majority (12, 57%) of these key AS events functionally linked to the phenotypic defects are exclusively identifiable by JUM. 1 (5%) of the AS events are identified by all three methods. 4 (19%) AS events are predicted by both rMATS and JUM and 4 (19%) AS events by MISO and JUM. See also Table S8. (B) Test if JUM can also identify rMATS-predicted, significantly changed AS events in genes associated with male courtship regulation in the PSI mutant Drosophila strain. The majority of these rMATS-predicted AS events are identified by JUM also (32, 80%), among which 67.5% of are re-classified correctly as Composite AS events by JUM (Figure S7A). 8 (20%) of rMATS-reported events are not identified by JUM, and later 3 (7.5%) events were found out to be incorrectly annotated AS events in the first place (Figure S7B, Table S9) and in 5 (12.5%) events the rMATS-annotated AS isoforms are either not expressed in the tissue sample or too lowly expressed to be detected by RNA-seq (Figure S7C, Table S9). (C) Number of significantly differentially spliced AS events reported in every AS pattern category by the three methods. (D) qRT-PCR validation of 12 significantly alternatively spliced AS events in genes associated with male courtship regulation that are only identified by JUM in the mutant male fly head. The Y-axis depicts the ratio between AS isoform a and isoform b, as indicated in the label below each bar graph. The detailed AS event structure and genome browser views of each AS event are provided in Figure S3-S6. The means of three independent measurements +/-standard deviation are shown. The differences between full-length WT PSI (blue) and PSI truncation mutation (PSIΔAB, red) male fly head samples were analyzed by oneway ANOVA test. (*) Statistically significant with P-value < 0.05; (**) statistically significant with P-value < 0.01; (***) Statistically significant with P-value < 0.001. (E) An example of an rMATS-predicted “SE” event that actually represents a much more complicated AS pattern in the male fly head and was re-classified correctly by JUM as a composite AS event is shown. Exon coverage from RNA-seq data is shown in blue; arcs represent splice junctions identified from the RNA-seq data; Drosophila annotation (dm3) of the transcripts is shown at the bottom. rMATS-predicted SE exon is specified with a red arrow. This SE exon is in fact alternatively spliced in combination with an upstream novel, unannotated alternatively spliced exon (marked by the single green arrow, whose existence was proven by the RNA-seq tracks and splice junction reads), as well as an A5SS site in the upstream exon (marked by double green arrow, and a zoom-in at that upstream exon is shown to provide a detailed view of the A5SS site).

For the counter test, we took the set of 40 rMATS-identified, significantly changed AS events that are within genes associated with male courtship behavior regulation and asked if JUM can identify these AS events as well (Figure 7B, Table S9). We found that among them, 5 events (12.5%) are identified by JUM also as significantly changed AS events in the category classified by rMATS (Figure 7B, Table S9); 27 events (67.5%), are identified by JUM also as differentially spliced AS events, but re-classified as composite AS events and a visual examination using the igv genome browser confirmed the predictions of JUM (Figure 7B, Figure S7A, Table S9). 8 events (20%) are not identified by JUM. However, when we examined these events individually on genome browser, we found that 3 events (7.5%) are incorrectly annotated AS events called by rMATS in the first place (Figure 7B, Figure S7B, Table S9). As for the rest, 5 events (12.5%) the rMATS reported/annotated AS isoforms are either not expressed or too poorly expressed to be detected by RNA-seq in the head tissue samples under study (Figure 7B, Figure S7C, Table S9). These results suggest that JUM is capable of identifying true differentially spliced AS events and profiling the events into the correct AS pattern category compared to other annotation-based software.

We further examined the distribution of differentially spliced AS events predicted by each method across different AS pattern categories (Figure 7C). rMATS again reported the highest number of differentially spliced AS events in SE (1716 out of a total of 3,634; 47%), and MISO reported the highest number of AS change in IR (1248 out of a total of 2,192; 57%). JUM, on the other hand, reported the highest number of change in A5SS (580 out of a total of 2245; 26%) and again reported a much less skewed distribution in the other AS pattern categories (217 in SE, 360 in A3SS, 183 in IR and 25 in MXE) than rMATS and MISO. Importantly, the changed AS event distribution reported by JUM correctly reflects the functional association of the specific PSI mutation with U1 snRNP and the regulation of 5’ splice site usage. Taken together, we conclude that JUM is not only able to accurately detect novel, tissue-specific AS events that are missed or mis-classified by other annotation-based methods, but JUM’s unique feature of accurately assembling AS patterns directly from RNA-seq data can be useful in predicting the functions of splicing regulators from the global AS changes caused by the regulator. Such unique features are not found in other currently available AS analysis methods.

## DISCUSSION

As a major mechanism for eukaryotic gene regulation, AS generates exceptionally diverse patterns of mRNA populations and their encoded proteins in metazoans. Different tissues, even sub-cellular populations within a given tissue or organ possess their own distinct AS profiles that are dynamically altered over temporal stages of development and cellular activities. The diversity and dynamics of AS patterns impose a major challenge for computational tools to quantify and compare AS profiles from RNA-seq data. Currently available AS analysis software tools commonly employ a top-down strategy based on pre-built annotated collections of known AS events to outline the general picture of splicing patterns for downstream analysis. This strategy greatly facilitates downstream quantitation, but at the same time fails to address the diversity in the patterns of AS, even when aided with workarounds to include novel splicing events specific to the sample under study. More importantly, the philosophy of having AS analysis depend on a pre-annotated AS event library or transcriptome that is usually compiled from all tissues and cell type mRNAs from the organism available in public databases underestimates the diversity and tissue-specificity of AS, which is often unique for a given tissue or cell type (Figure S1), and can be vastly different from the prior known annotations or will only identify a subset of the annotated AS patterns. On the other hand, the top-down strategy restricts the quality of downstream AS analysis depending on how extensive the annotation is. Technically speaking a complete, absolutely comprehensive annotation of all mRNA types in metazoan tissues can be achieved over time by deeply sequencing all possible tissues and cell subtypes. However, the reality is that the limits of complete transcriptome annotation require great effort to reach and so far, the annotation for known AS events is far from complete, and is especially poor when it comes to tissues with extraordinarily complex AS patterns, such as neurons, gonads and cancer samples, or in organisms other than human and mouse.

With JUM, we approach the problem of tissue-specific AS pattern analysis with a different philosophy, the bottom-up approach that profiles, quantitates and analyzes AS patterns directly from the sample under study (Figure 1 and Figure 2). By utilizing the unique topological features of the splicing graph representing each AS pattern, JUM is able to accurately construct the sample-specific AS atlas through assembling the basic graphical nodes of the atlas called AS structures that are profiled directly from the sample. JUM then performs differential AS analysis through building robust statistical power from quantifying raw read counts spanning splice junctions in AS structures (Figure 1). The approach that JUM uses not only provides a thorough and statistically robust investigation of the diverse and dynamic AS patterns specific to a given biological sample, but also eliminates the labor-intensive computational effort required to build extensive AS event annotations or transcriptome assembly. It needs to be noted that although JUM is independent of transcriptome annotation, it is capable of accurately profiling AS events that are previously known. In fact, a significant percentage of the JUM output results cover previously annotated AS events (Figure 6B and Figure 6D). It is also worth noting that although JUM relies exclusively on splice junction reads from the specific sample and profiles tissue-specific AS patterns that are novel, these JUM-identified AS events are not necessarily in poorly annotated or lowly expressed genes in the tissue sample. For example, the male courtship behavior-associated differentially spliced AS events that are only identified by JUM in PSI mutant male fly heads are in genes are that abundantly expressed, with FPKM ranges from 20 to 200 (Figure 7A, Figure S3-S6). It should also be noted that a recent method called LeafCutter also performs splicing analysis independent of transcriptome annotation (21). However, LeafCutter emphasizes only quantifying levels of intron excision, without regard to detection, quantification or analysis of AS event patterns. LeafCutter is also a tool more specialized for analyzing eQTLs (21).

Another novel feature of JUM lies in its stringent statistical platform to analyze IR events. IR is a crucial AS regulatory mechanism for gene control but the complex composition and structure of IR pattern makes it easy to misidentify other AS types as IR events, causing a high false positive rate of IR event profiling and quantification (Figure 3). Thus, JUM offers a new and well-developed approach to analyze IR specifically, which uses three carefully designed standards that filter out most of the AS types that can be easily mistakenly classified as IR by conventional methods in currently available AS analysis tools (Figure 3). As a result, JUM provides a reliable means to accurately evaluate the importance of IR in cellular activities and diseased cellular states (Figure 5).

A third unique feature of JUM is that it classifies a separate AS pattern category which is not usually covered in other currently available AS methods: the Composite AS events. This is a class of AS event that is an inseparable combination of the conventionally recognized AS patterns and cannot be simply categorized and quantified as one of the decomposed AS patterns. Composite AS events are widespread and found in tissue types with diverse AS profiles, such as neurons and cancer cells, and their existence directly reflects the complexity of AS and these novel isoforms can play important roles in shaping cellular activities and physiology. JUM thus offers a way to profile and classify this specific class of AS patterns and will facilitate the investigation of this class of previously under-studied AS events.

Last but not the least, JUM possesses the unique feature of accurately profiling the changed AS events in terms of the standard AS pattern categories of SE, A5SS, A3SS, IR and MXE directly from the RNA-seq datasets. This feature is important but not commonly achieved in other currently available AS analysis tools, especially in tools that rely less on pre-annotation of the transcriptome. For example, LeafCutter can analyze the alternative excision of introns without dependence on annotation, but it does not differentiate AS changes in terms of splicing patterns (21). For annotation-based computational tools that report AS analysis results in the different AS pattern categories, bias towards one type of pattern is consistently observed (see above). For example, MISO tends to bias towards SE and IR events and rMATS tends to bias towards SE events (Figure 6, Figure 7, Table S5 and Table S6). The ability to profile AS changes into AS pattern categories is especially important when investigating global AS changes brought about by the perturbation of a splicing regulator. As splicing regulators affect different steps of intron splicing, such as choices of 5’ splice site, 3’ splice site or branch point, the regulatory mechanism of a splicing regulator can be reflected in the distribution of changed AS events among the different AS pattern categories. As an example, the splicing factor PSI in Drosophila regulates 5’ splice sites through its interaction with U1 snRNP (47). Using JUM, we found that indeed A5SS is the most affected splicing pattern in the male head transcriptome of a Drosophila strain that carries a mutation in PSI that disrupts its interaction of U1 snRNP. In this scenario, the accurate classification of splicing changes in AS patterns by JUM predicts the regulatory mechanism of PSI in AS regulation. Thus, JUM not only offers a comprehensive and accurate method for global AS analysis, but also provides a framework to indicate the regulatory function of splicing factors that cause the global AS changes.

Multiple applications of JUM in analyzing complex tissue types consistently showed that JUM is capable of detecting true, novel and unannotated AS changes that are not detected by other annotation-based methods. Previously, we used earlier versions of JUM to analyze two sets of neuronal RNA-seq datasets, one prepared from mouse embryonic cortical neurons experiencing a dendrite outgrowth defect upon the knockdown of the splicing regulator PQBP1 (48), and one from the male brain of a Drosophila strain whose males suffer from aberrant courtship behavior due to mutation of the splicing regulator PSI (46). We asked if JUM could identify significantly changed true AS events that can be associated with the phenotypic defects at the molecular level in the neuron cell type that present the most diversified, dynamic, novel and unannotated AS patterns (1, 5). We found that JUM identifies a set of key neuronal AS events in these samples that are functionally linked to the phenotype. We used RT-PCR to verify this set of JUM-predicted AS events in mouse and Drosophila samples and validated the AS changes for all of them (46, 48). Importantly, many of these experimentally validated, crucial AS events were only identified using JUM, but not using other AS analysis tools in comparison (46, 48). For example, the significant AS changes of the Drosophila *fruitless* mRNA transcript isoforms that are directly linked to abnormal male courtship behavior in mutant PSI Drosophila males were captured exclusively by JUM, but not MISO (10) or JuncBase (14). In fact neither the AS annotations used by MISO nor the *de novo* assembled transcriptome annotation supplied to JuncBase included the AS of the last exons of the *fruitless* male transcripts in the first place (Table S8). The other example is that the significant AS changes of the *Ncam1* transcripts that are functionally associated to the dendritic outgrowth defect in PQBP1-perturbed mouse neurons were discovered only by JUM, but not DEXSeq (11, 48). Even when DEXSeq was supplied with a *de novo* assembled transcriptome annotation that included the alternatively spliced cassette exon in *Ncam1* transcripts, DEXSeq was not able to detect the significant changes, possibly because of the small size of the exon (30bp) (48). These results further suggest that JUM can identify functionally important tissue-specific AS events with accuracy and specificity.

JUM exclusively use reads mapped over splice junctions for downstream quantification, and thus prefers that RNA-seq libraries be sequenced at a deeper level (typically duplicates or triplicates of > 40-50 million Illumina reads for human cell lines), compared to other tools that also utilize exonic reads. We feel that with the cost of sequencing dropping rapidly, deeper sequencing is a small price to pay to comprehensively investigate tissue-specific AS patterns. Exonic reads may help increase the statistical power when quantifying differential AS changes, however it requires the input of transcriptome annotations. Since splice junction reads are the most direct evidence for the quantitative assessment of splicing, with a well-sequenced RNA-seq dataset splice junction reads should be prevalent enough to quantitate AS changes without the need for exonic reads.

In conclusion, JUM presents a novel and statistically rigorous approach to address, evaluate, quantitate and classify the complex and diverse patterns of AS profiles in eukaryotic transcriptomes. We are confident that this new approach will provide new and important insights to the dynamic regulation of AS and gene expression. Our results indicate superior performance of JUM in novel AS detection, quantification and classification from Drosophila heads, mouse neurons, human cancer tumor and human cancer-associated splicing factor mutation carrying cell line RNA samples. These initial applications already indicate that JUM will be the method of choice going forward to analyze complex pre-mRNA splicing patterns at the transcriptome-wide level, particularly in complex cell or tissue types that are already known to generate extremely diverse mRNA isoform profiles, such as gonads (testes and ovaries), pluripotent stem cells and a variety of neuronal cell types and nervous system tissues. JUM can also be readily extended to analyze AS patterns in single cells. Finally, the JUM algorithm should be useful for detecting, mapping and analyzing the biogenesis of a novel pattern of non-coding RNAs that are circular in structure. These newly discovered RNA species are thought to play a role in microRNA regulation (49).

## MATERIALS AND METHODS

### RNA-seq data

Raw RNA-seq data (FASTQ format) for Drosophila male fly heads and K562 cell lines with SRSF2 mutations described in the paper are derived as previously described (45, 46). Human colon tumor and matched normal tissue poly-A selected RNA-seq data (in BAM format) are acquired from the TCGA database. A detailed description of the patient tumor and normal samples used in this study are listed in Table S4. The downloaded BAM files are transformed back to FASTQ format by using the SamToFastq function in PICARD tools before analysis. The FASTQ data are then mapped to the human genome hg38 as described below. The sequencing read mapping results are summarized in Table S7 for each patient.

### RNA-seq data mapping for JUM

RNA-seq reads are mapped to the human (hg38) (or hg19 for MISO as the pre-annotation of splicing events provided by MISO is in hg19) and Drosophila (dm3) genomes respectively using STAR (50) in the 2-pass mode, as instructed in the STAR manual. Only unique mapped reads are kept in the output for downstream JUM analysis.

### RNA-seq data experiment simulation

We used the ASmethodBenchmarking software as described in (42) to simulate RNA-seq datasets, triplicates of ~80 million 100bp reads are simulated for each condition, with three levels of AS changes. Parameters are listed in Table S1.

### Running MISO, MAJIQ, Cufflinks and rMATS on simulated RNA-seq datasets for differential AS analysis

The detailed commands for each software as well as the versions of each software used in this study are listed in Table S2.

### ROC curve potting and AUC calculation

Ranking score for ROC curve plotting is chosen to be 1-adjusted_pvalue for JUM, Cufflinks and rMATS, as the three methods provide adjusted p-values from multiple testing correction. For MAJIQ and MISO that do not offer multiple testing, the ranking score is set to be the maximum value of E(dPSI) among LSV junctions in an AS event for MAJIQ (18) and the value of Bayes factor for MISO (10). The R package ROCR (51) is used to plot ROC curves and calculating AUC metric.

### Algorithm to construct AS patterns from profiled AS structures

We first profile all AS structures from the RNA-seq data and calculate the *S_I_* value for each sub-AS-junction in these AS structures. Two AS structures are defined as “linked”, if they share one specific sub-AS-junction, and a “path” is drawn between the two AS structures. Under this definition, a “loop” of AS structures are searched in the whole pool of AS structures, with every AS structure in the loop linked to one other by a path. Each profiled loop of AS structures is corresponding to an AS pattern, and is allocated to each AS pattern category based on the features of the sub-AS-junction *S_I_* value distributions.

### Visualization

All RNA-seq track data and junction reads are visualized using IGV (44) and the Sashimi plots tool (52).

### RNA extraction and qRT-PCR validation of JUM-predicted AS events

Drosophila heads were isolated from 10-20 manually sorted and snap-frozen males from the PSI mutant and wildtype PSI strains as previously described (46). RNA were extracted using the Trizol reagent. qRT-PCR primers were designed with the software Primer3 (http://bioinfo.ut.ee/primer3-0.4.0/) and qRT-PCR experiments performed by using the SuperScript III Platinum SYBR Green One-Step qRT-PCR Kit (Thermo Fisher Scientific) on a LightCycler 480 instrument (Roche). All AS events were validated as described above except for Fmr1 and lig gene AS events, as no designed primers produced one single product for qRT-PCR experiments. Thus, for these two genes RNA (0.5 μg) were first reverse-transcribed by SuperScript First-Strand Synthesis System (Invitrogen) and the splicing isoforms were analyzed by RT-PCR followed by gel electrophoresis on 6% TBE gel (Thermo Fisher Scientific), stained with SYBR Gold (Invitrogen) and quantified with ImageJ software (NIH).

### Parameters used for AS analysis tool comparison

A detailed command list for running rMATS, MISO and IRFinder described in this study is listed in Table S2 and Table S3.

For colon cancer patient data analysis, we used the following statistical cutoff:

MISO: at least 10 unique reads mapped to each isoform, splicing change at least 10%, Bayes factor 5.

rMATS: adjusted pvalue <= 0.05.

JUM: adjusted pvalue <= 0.05, splicing change at least 10%

IRFinder: pvalue <= 0.05, splicing change at least 10%

For SRSF2 mutation carrying K562 cell line analysis, we used the following statistical cutoff:

rMATS: adjusted pvalue <= 0.1, splicing change at least 10%

JUM: adjusted pvalue <= 0.1, splicing change at least 10%.

For Drosophila male head samples that carry a PSI mutation analysis, we used the following statistical cutoff:

MISO: at least 5 unique reads mapped to each isoform, splicing change at least 5%, Bayes factor 5.

rMATS: adjusted pvalue <= 0.1, splcing change at least 5%.

JUM: adjusted pvalue <= 0.1, splicing change at least 5%.

### Software availability

A user-friendly version of the JUM package has been deposited on GitHub https://github.com/qqwang-berkeley/JUM. The codes are written in perl and bash shell scripts.

## Acknowledgements

We thank Jeffrey Paulsen, Chao Di and Kate Abruzzi for helpful critiques and comments. We thank Yeon Lee for help testing the first user-friendly version of the JUM package. We thank Ashley Albright and the Michael Eisen lab for help with the qRT-PCR experiments. The results shown here are in part based upon data generated by the TCGA Research Network: http://cancergenome.nih.gov/. This work was supported by NIH R01GM097352 and NIH R35GM118121 (D. Rio, PI) and by the NIH Center for RNA Systems Biology at U.C., Berkeley (P50GM102706; J. Cate, PI). Q.W. is supported by the Arnold O. Beckman Postdoctoral Fellowship.

## Disclosure declaration

The authors declare no conflicts of interest.

